# *Measuring* Clinically Relevant Knee Motions With A Self-Calibrated Wearable Sensor

**DOI:** 10.1101/309328

**Authors:** Todd J. Hullfish, Feini Qu, Brendan D. Stoeckl, Peter M. Gebhard, Robert L. Mauck, Josh R. Baxter

## Abstract

Low-cost sensors provide a unique opportunity to continuously monitor patient progress during rehabilitation; however, these sensors have yet to demonstrate the fidelity and lack the calibration paradigms necessary to be viable tools for clinical research. Therefore, the purpose of this study was to validate a low-cost wearable sensor that accurately measured peak knee extension during clinical exercises and needed no additional equipment for calibration. Knee flexion was quantified using a 9-axis motion sensor and directly compared to motion capture data. Peak extension values during seated knee extensions were accurate within 5 degrees across all subjects (RMS error: 2.6 degrees, *P* = 0.29) but less accurate during sit-to-stand exercises (RMS error: 16.6 degrees, *P* = 0.48). Knee flexion during gait strongly correlated (0.84 ≤ r_xy_ ≤ 0.99) with motion capture measurements but demonstrated average errors of 10 degrees. This study demonstrated a low-cost sensor that satisfied our criteria: a simple calibration procedure resulting in accurate measures of joint function during clinical exercises, making it a feasible tool for continuous patient monitoring to guide regenerative rehabilitation.

## Introduction

Post-operative knee stiffness following total knee arthroplasty often leads to flexion contracture deformities that require aggressive therapies and revision surgery [1]. Restoring knee motion within three months following joint replacement surgery mitigates the risk of flexion contracture and poor outcomes [2,3]. While motion capture accurately quantifies knee flexion [4], such measurements are financially and logistically impractical for continuously monitoring post-operative patient progress. Outpatient physical therapy remains the most common means of post-operative care, but this approach is costly, time-consuming, and ultimately depends on patient compliance. As such, there is a need for cost-effective means of remote, continuous tracking of patient knee motion following invasive surgery.

Therefore, the purpose of this study was to validate a novel implementation of a low-cost wearable sensor [In Review, proof attached] by satisfying several criteria: 1) simple calibration procedure, 2) accurate measurement of peak knee extension during clinical exercises, and 3) quantification of knee motion during walking. To that end, we compared the knee angle measurements derived from the wearable sensor to those calculated by motion capture during clinically relevant functional activities. Based on clinician input, we established a clinically acceptable peak-knee extension error of less than 5° during a clinical exercise compared to measurements using motion capture.

## Methods

Seven healthy young adults (4 males, 3 females; 26 ± 4 years; BMI 23.8 ± 3.7) participated in this IRB approved study to establish the feasibility and accuracy of a low-cost wearable sensor. Subjects wore standardized attire and reflective markers placed on the lower-extremities were tracked using a 12-camera motion capture system (Raptor Series, Motion Analysis Corp, Santa Rosa, CA). Marker trajectories acquired during a standing trial were used to scale a constrained-kinematic lower-extremity model [5]. Subjects performed clinical motions – seated knee extension and sit to stand – and walked on a treadmill for 2 minutes at three speeds (0.9, 1.2, and 1.5 m/s) and up a 10% grade (1.2 m/s).

Knee motion was quantified using a wearable magnet-based device [manuscript in revision] that was positioned on the right leg to measure knee flexion during functional activities. The magnetic field strength of a rare earth magnet (1.50”x0.50”x0.25” Neodymium Block Magnet, CMS Magnetics, Garland, TX) was measured using 9 degree-of-freedom inertial measurement unit (IMU, LSM9DS0, FLORA 9-DOF, Adafruit, New York, NY). The magnet and IMU were secured to the distal-lateral thigh and proximal-lateral shank, respectively, using fabric-backed tape and self-adhesive wrap. The accelerometer, gyroscope, and magnetometer ranges were set to ± 2g, ± 245°s^−1^, and 12 gauss, respectively. Sensor data were synchronized with the motion capture system via a digital pin on the microcontroller (FLORA – Wearable Platform, Adafruit, New York, NY) and logged to file.

The wearable sensor was calibrated by modeling flexion angle as a function of the magnetic field strength during a series of static knee postures. Subjects sat on a treatment table with the thigh parallel to the tabletop and lower-leg freely hanging in approximately 90 degrees knee flexion.

Next, the knee was held at five evenly spaced positions between 90 and 0 degrees knee flexion for several seconds (**Fig. 1**) as the magnetic field strength and direction of gravity were acquired using the magnetometer and accelerometer, respectively. Subjects then stood with the lower leg vertically aligned to the ground to account for IMU orientation on the shank. Next, flexion angle was approximated using the lower leg pitch and used to fit a 3^rd^ order polynomial to the magnetic field strength. Lower-leg angular velocities were collected during the quiet standing trial to tare the gyroscope signal. Marker trajectories and IMU data were collected during each functional activity.

**Fig. 1:**
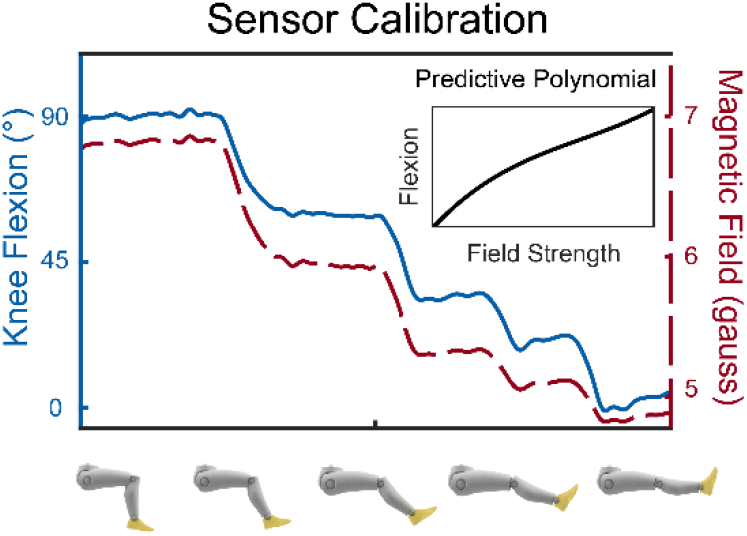
Sensor calibration was performed by placing the knee in 5 arbitrary positions and fitting a 3^rd^ order polynomial (inset) to the magnetometer-accelerometer data.

Lower extremity kinematics were calculated using a constrained-kinematic model (Opensim v3.3; [6,7]). Marker trajectories and the wearable sensor measurements were filtered using a 6 Hz fourth-order Butterworth filter. Heel strike events during gait were identified using a kinematic-based algorithm [8]. Knee flexion angles and tibial angular velocities were calculated for each stride and averaged across approximately 80 strides per subject. Angular velocities of the lower-leg were also calculated from segmental measurements calculated from the constrained-kinematic model and gyroscope.

Wearable sensor measurements were compared to motion capture measurements. The primary outcome was peak knee extension during a seated-knee extension with differences between measurement techniques of less than 5 degrees considered clinically acceptable. Knee angle and tibial angular velocity ninety-five percent confidence intervals were calculated for each movement using a bootstrap approach [9]. Peak knee extension and flexion during these activities were directly compared using paired t-tests (*p* < 0.05). Kinematic patterns were compared using cross correlation analyses to quantify the similarities between the wearable sensor and motion capture measurements [10].

## Results

Seated-knee extension was accurate to within 5 degrees across all subjects (*P* = 0.29; RMS error: 2.6 degrees; Fig. 2A). Peak knee extension measurements were less accurate during the sit to stand exercises, consistently under-approximating extension values (*P* = 0.48; RMS Error: 16.6 degrees, Fig. 2B). Peak knee flexion during both of these movements reached sensor saturation at approximately 65 degrees knee flexion (**Fig. 2**).

**Fig. 2:**
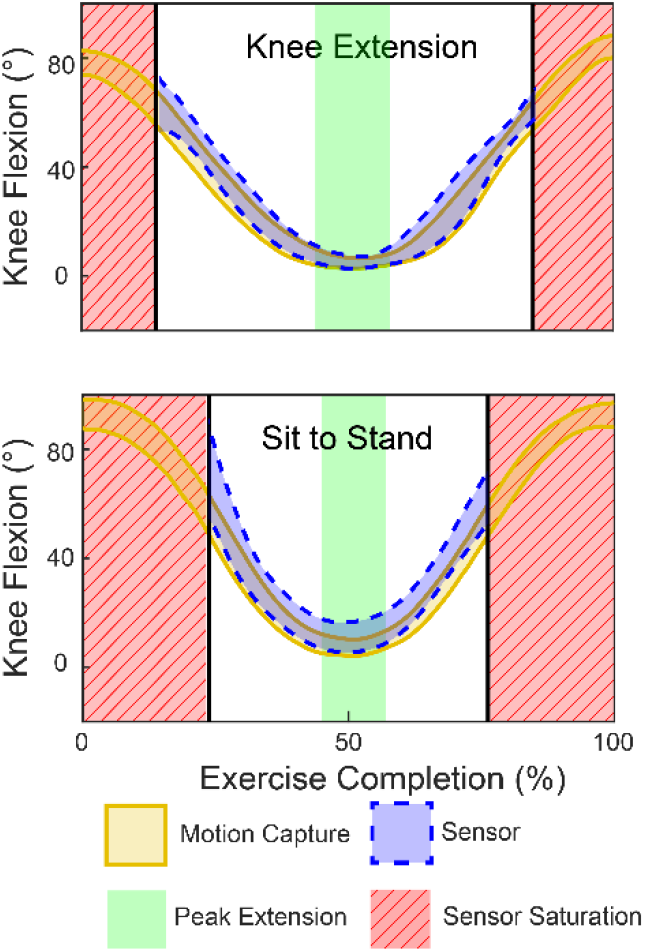
Knee angles measured by the device (dashed blue) were compared to motion capture measurements (solid gold) across a single extension event. Peak knee extension (green area) did not differ between the two measurements. The sensor became saturated with increased knee flexion (red stripped area).

Knee flexion angles and tibial angular velocities strongly correlated with motion capture measurements during walking (0.84 ≤ r_xy_ ≤ 0.99; **Fig. 3**). Despite similarities in kinematic patterns, the wearable sensor showed an RMS Error of 10.3 degrees peak knee flexion. Walking faster and up an incline generated smaller errors (*P* = 0.47; RMS errors: 7.9 and 7.4 degrees, respectively) compared to walking at slow and medium speeds generated (*P* = 0.64; RMS errors: 12.8 and 13.2 degrees, respectively). Tibial angular velocities measured using the gyroscope detected 22.1 degrees/s increases with 0.3 m/s increases in walking speed (*P* < 0.05).

**Fig. 3:**
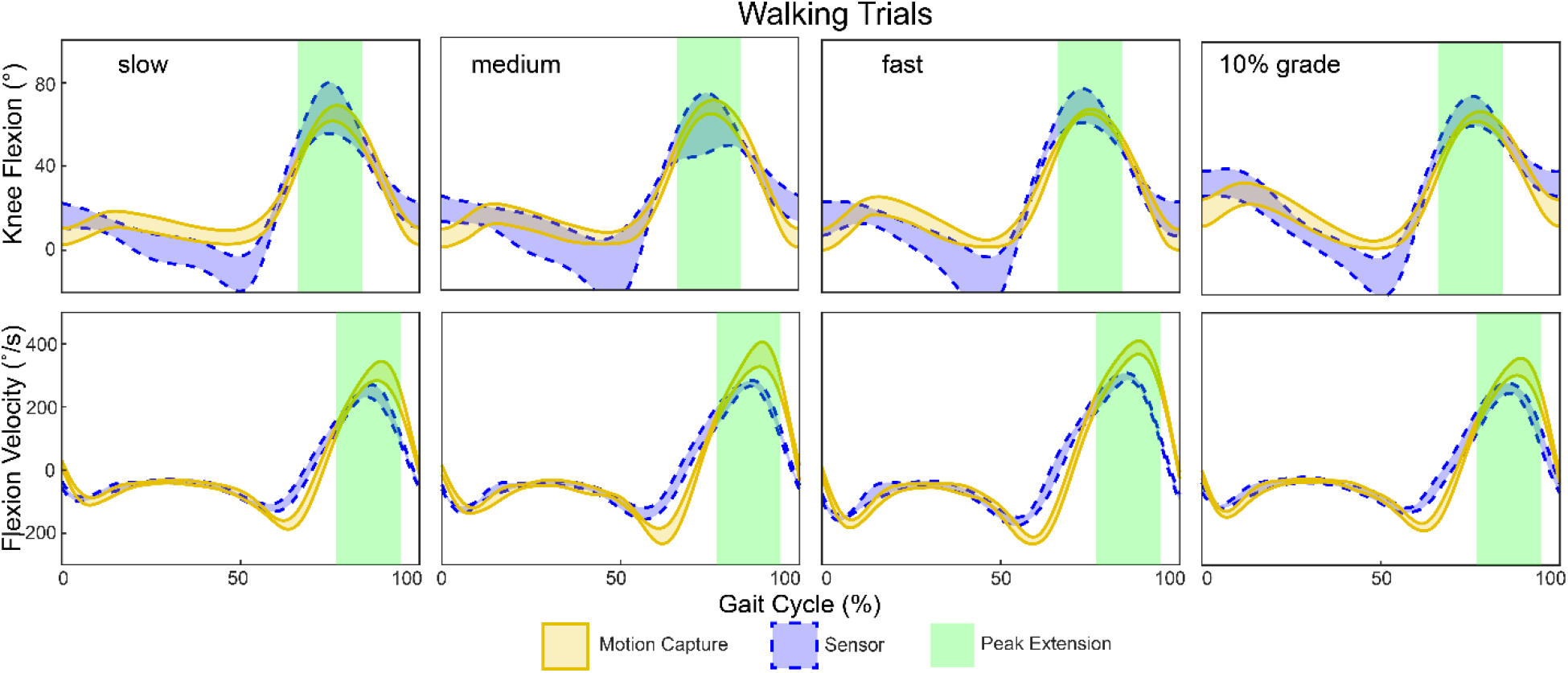
Knee angles and tibial angular velocities strongly agreed (r_xy_ ≥ 0.84) between the device (dashed blue) and motion capture (solid gold). In addition, peak knee flexion measurements were compared at peak knee flexion (green area).

## Discussion

This study tested a new paradigm for measuring clinically relevant joint motions using a single low-cost sensor that does not need additional equipment to calibrate. Our results support the feasibility of this sensor paradigm for measuring knee extension during a seated-knee extension (**Fig. 2**). However, more dynamic motions like walking appear to be affected by soft tissue artifact and sensor limitations, which produce less accurate predictions of knee angle (**Fig. 3**). While this study focused on measuring knee motion, this paradigm could be adapted to work with other mostly-planar joints such as the elbow or ankle. Monitoring knee motion using a low-cost sensor provides new opportunities for providers to monitor patient progress and function outside of clinical visits.

This study was affected by several limitations. First, only healthy young adults participated in this study to establish the feasibility of the sensor and calibration paradigm. These subjects had a lower body mass index (23.8 ± 3.7) than many patient populations, including those undergoing total knee arthroplasty [11] and a single investigator placed the sensor on all research subjects. Soft-tissue artifact around the knee joint [4] likely affected our measurements and other technical paradigms may be less sensitive to skin motion [12,13].

Low-cost wearable sensors offer an exciting alternative to traditional motion capture techniques [12,13]. In the present study, we present a unique implementation of an IMU in conjunction with a magnet to accurately measure clinically-relevant knee motion. Low-cost sensors may address concerns of increasing costs and decreasing accessibility in healthcare. Establishing predictors of patient function and outcomes with low-cost wearable sensors may help detect adverse events, guide regenerative rehabilitation, and improve patient-specific treatments.

## Acknowledgments

We thank Ms. Annelise Slater for assistance with data collection.

## Notes

***Funding*** no funding has been provided for this research

